# A Shared Neural Marker Predicts Creative Performance Across Distinct Problem-Solving Tasks

**DOI:** 10.1101/2025.09.16.676681

**Authors:** Zenas C. Chao, Miyoko Street, Tzu-Ling Liu

## Abstract

Creativity is essential for innovation, yet the brain mechanisms supporting its moment-to-moment variability remain unclear. We hypothesize that creativity depends on dynamic fluctuations in neural flexibility, which determine the potential to generate creative solutions. Here, we identify a shared neural marker of “creativity potential” that predicts upcoming performance across distinct problem-solving tasks. Twenty-eight participants completed the Alternative Uses Test (AUT), a measure of divergent thinking, and the Fusion Innovation Test (FIT), which integrates divergent and convergent thinking. Responses were scored for novelty, feasibility, and goal attainment using validated GPT-based automated evaluation. EEG signals recorded prior to problem onset were used to decode single-trial creativity scores. A decoding model based on coherence features achieved robust performance (*r* = 0.45, leave-one-trial-out) and generalized across individuals (*r* = 0.34, leave-one-subject-out). Feature weights revealed a creativity potential network (CPN), characterized by frontal-temporal interactions in beta frequency band. Applying the model to resting-state recordings revealed ∼3-minute cycles of creativity potential, suggesting intrinsic brain dynamics shape readiness for creative problem-solving. These findings establish a shared neural marker of creativity that transcends task boundaries and individuals. Beyond advancing our understanding of creative cognition, this work opens the possibility of monitoring creativity potential in real time, with implications for neurofeedback and creativity enhancement in daily life.

**Significance statement:** Creativity allows us to generate novel and useful ideas, yet our ability to be creative fluctuates from moment to moment. Here we identify a neural marker of “creativity potential” that predicts upcoming creative performance across two distinct problem-solving tasks. Using EEG and GPT-based automated evaluation, we show that preparatory brain activity encodes creativity potential, generalizing across tasks and individuals. Furthermore, creativity potential fluctuates in intrinsic ∼3-minute cycles during rest. These results advance our understanding of the neural basis of creativity and provide a foundation for real-time monitoring and neurofeedback applications that may help individuals enhance their creative capacity.

## Introduction

During problem solving, sometimes solutions emerge easily, while at other times they remain elusive. We hypothesize that such variability reflects fluctuations in the brain’s flexibility, which determine the potential to generate creative solutions. We refer to this as “creativity potential.” Converging evidence suggests that creativity depends on flexible brain states that allow dynamic reconfiguration of neural networks (1–5). For example, in the Remote Associates Test, a convergent thinking task, brain activity preceding problem presentation differs between trials that do and do not evoke an “Aha” experience (6). However, reliance on subjective insight is problematic, as “Aha” experiences can accompany incorrect solutions (7) and even bias creativity evaluation (8). These findings highlight the need for more objective, task-general measures of creative performance.

To address this, we employed complementary assessments of divergent and convergent thinking. The Alternative Uses Test (AUT) asks participants to generate unconventional uses for everyday items (e.g., umbrella) (9). Responses are evaluated for novelty and feasibility using a validated GPT-based method that matches human judgments (10). While the AUT captures divergent idea generation, real-world creativity also requires narrowing down to effective solutions. To capture this process, we used the Fusion Innovation Test (FIT), which requires participants to solve open-ended problems related to personal or societal goals under the constraint of combining unrelated elements. The FIT evaluates both divergent and convergent processes, with responses scored for novelty, feasibility, and goal attainment using automated GPT-based evaluation (11).

Using EEG, we asked whether a shared neural marker predicts creative performance across both tasks. Specifically, we focused on brain activity preceding problem onset, reasoning that preparatory brain states encode creativity potential. With both leave-one-trial-out and leave-one-subject-out cross-validation, our decoding model achieved robust prediction of creativity scores (*r* = 0.45 and 0.34, respectively), demonstrating generalizability across tasks and individuals. Furthermore, applying the model to resting-state recordings revealed ∼3-minute cycles in predicted creativity potential, suggesting intrinsic fluctuations in the brain’s readiness for creative problem solving. By identifying a neural marker of creativity potential, this work provides a foundation for monitoring creativity in everyday life and developing real-time neurofeedback to enhance creative ability.

## Results

### Creativity tasks and overall creativity assessment

We examined creativity using two established tasks: AUT and FIT, with example questions and solutions shown in Figures 1A and 1B. Data were collected from 28 participants (17 males and 11 females; age range 18∼29 years, mean ± standard deviation: 22 ± 2.5 years). Each participant completed 30 AUT questions and 30 FIT questions, administered across two experimental days. On each day, participants performed one block of 15 AUT questions and one block of 15 FIT questions. The order of task blocks was reversed on the second day, and the starting task (AUT or FIT) was counterbalanced across participants. After both sessions, participants also completed an online set of questionnaires to assess their everyday creativity.

**Figure 1.**
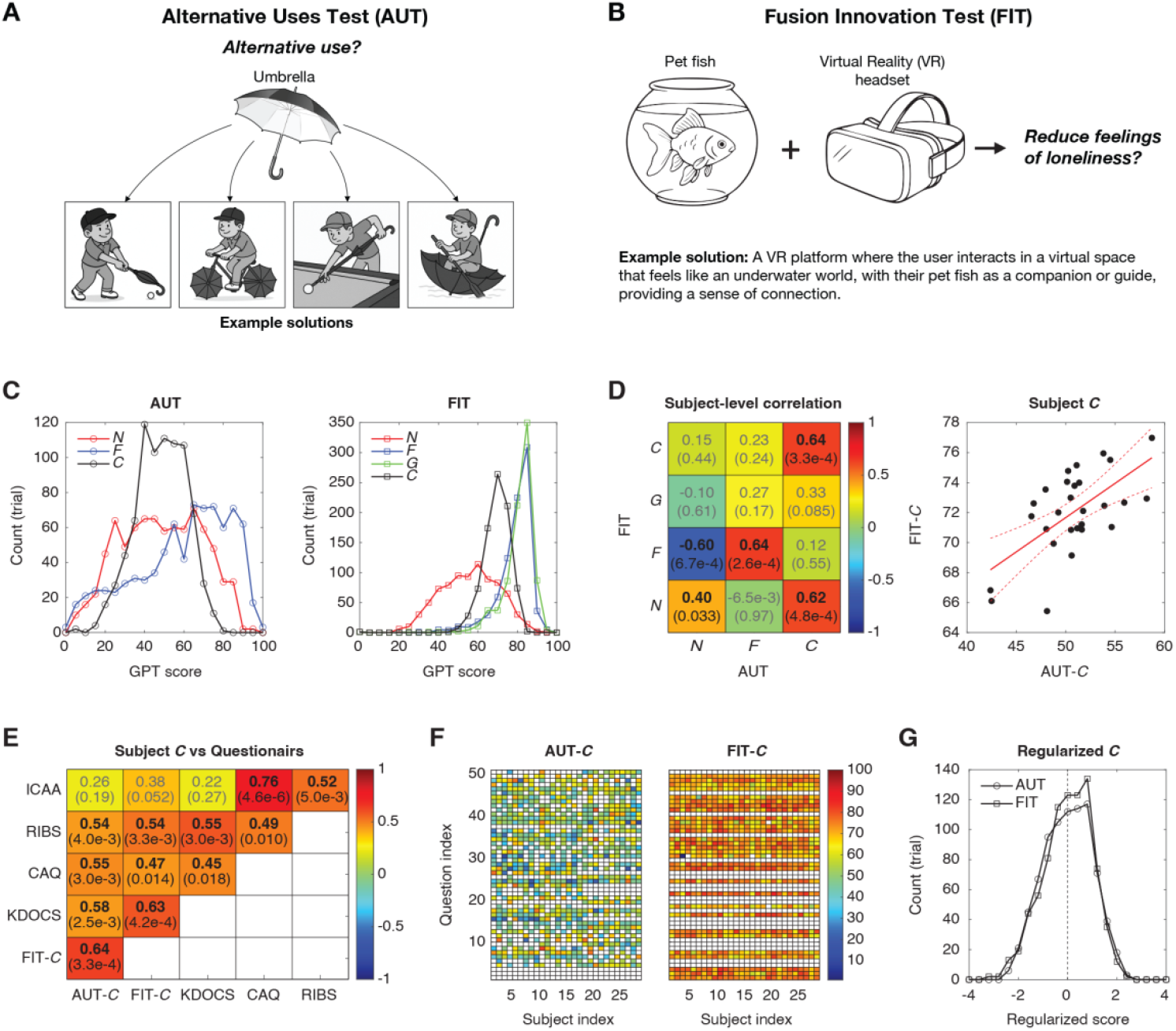
Creativity tasks and behavioral analysis. (**A**) Alternative Uses Test (AUT). The example item shown is “umbrella,” with possible creative solutions illustrated. In the actual test, both items and responses are presented in text only; figures here are for illustration. (**B**) Fusion Innovation Test (FIT). Two sample question items and a goal are presented, along with an example solution. As with AUT, the real test uses text only. (**C**) Histograms of GPT-based scores for novelty (*N*), feasibility (*F*), and overall creativity (*C*) in AUT (left), and for novelty (*N*), feasibility (*F*), goal attainment (*G*), and overall creativity (*C*) in FIT (right), across all solutions from all participants. (**D**) Subject-level correlations between AUT scores (x-axis) and FIT scores (y-axis). Colors indicate correlation coefficients, with correlation coefficients and p-values (in parenthesis) labeled. Significant correlations are shown in bold black, non-significant ones in gray. The correlation between subject-level creativity scores (AUT-*C* vs. FIT-*C*) is shown on the right. Each black circle represents a subject’s mean score. Red line: linear regression; dashed lines: 95% confidence interval. (**E**) Correlations between subject-level AUT-*C*, FIT-*C*, and four questionnaires (KDOCS, CAQ, RIBCS, ICAA), using the same format as in panel D. (**F**) Heatmap of AUT-*C* and FIT-*C* by subject and question (note: not all questions were used). Color indicates score. (**G**) Histograms of regularized creativity scores (*C*) for AUT and FIT. Images in panels A and B were generated with ChatGPT (OpenAI).

AUT responses were scored for novelty (*N*) and feasibility (*F*) using a GPT-based automated evaluation method (see details in Methods) (10). Overall creativity (*C*) was defined as the geometric mean of *N* and *F* (*C* = √(*N* × *F*)). Distributions of *N, F*, and *C* across the 840 AUT trials (28 participants × 30 questions) are shown in Figure 1C (left). FIT responses were evaluated for novelty (*N*), feasibility (*F*), and goal attainment (*G*) using a similar GPT-based method (see details in Methods) (11). Overall creativity (*C*) was defined as the geometric mean of *N, F*, and *G* (*C* = ^3^√(*N* × *F* × *G*)). The distributions of *N, F, G*, and *C* from the 840 FIT trials are shown in Figure 1C (right).

Note that AUT feasibility (AUT-*F*) and FIT feasibility (FIT-*F*) and goal attainment (FIT-*G*) scores were skewed toward higher values, while novelty scores (AUT-*N* and FIT-*N*) were more symmetrically distributed. Consequently, FIT overall creativity (FIT-*C*) was more strongly skewed toward higher scores compared to AUT overall creativity (AUT-*C*). This likely reflects the task constraints: in AUT, participants were required to propose alternative “uses” that balanced novelty and usefulness, making highly novel but impractical ideas (e.g., using an umbrella as a bicycle wheel; see Figure 1A) relatively rare. Similarly, in FIT, solving the problem imposed constraints on goal attainment and feasibility, prioritizing viable solutions over purely novel ones.

At the subject level, where scores were averaged across questions, correlations between AUT and FIT are shown in Figure 1D. Significant Pearson’s correlations were found for both novelty (*N*) and feasibility (*F*), indicating that generating creative uses of items relies on overlapping processes across the two tasks. In addition, strong cross-dimensional correlations emerged between AUT-*N* and FIT-*F*, as well as between AUT-*C* and FIT-*N*. Most importantly, overall creativity scores (AUT-*C* and FIT-*C*) showed the strongest correlation, suggesting that both tasks capture a shared dimension of individual creative ability. This subject-level correlation of overall creativity is further illustrated in Figure 1D (right).

To further validate the geometric mean as a measure of overall creativity, we examined the correlations of subject-level AUT-*C* and FIT-*C* with four established creativity-related questionnaires (see Methods for details). These included: the Kaufman Domains of Creativity Scale (KDOCS), which measures self-perceived creativity across multiple domains (12); the Creative Achievement Questionnaire (CAQ), which assesses real-world accomplishments in diverse areas (13); the Runco Ideational Behavior Scale (RIBS), which indexes creativity potential through the frequency of novel idea generation (14); and the Inventory of Creative Activities and Achievements (ICAA), which captures both creative activities and achievements across domains (15). As shown in Figure 1E, both AUT-*C* and FIT-*C* were significantly correlated with KDOCS, CAQ, and RIBS, and showed marginal correlations with ICAA, supporting their validity as indices of overall creativity.

For decoding purposes, AUT-*C* and FIT-*C* were regularized to account for distributional differences, particularly the skewness of FIT-*C* (Figure 1C). The original values of AUT-*C* and FIT-*C* are shown in Figure 1F, where the generally higher values of FIT-*C* are evident. To normalize these measures, we first calculated the z-score within each question, and then applied a second z-scoring across scores within each subject (see Methods for details and rationale). The resulting distributions of regularized *C* are shown in Figure 1G, where both AUT and FIT exhibit symmetric and comparable patterns. Values above zero indicate personally creative ideas, whereas values below zero indicate personally non-creative ideas.

### Creativity potential: decoding upcoming creative performance

During the task, we recorded 19-channel EEG signals using a wireless dry EEG system (Figure 2A; see Methods). The goal of decoding was to predict single-trial regularized *C* (from either AUT or FIT) based on pre-problem-solving brain activity. We refer to this as “creativity potential,” as it reflects the neural readiness for upcoming creative performance. To estimate this, we computed coherence across five frequency bands (delta, theta, alpha, beta, gamma) using EEG segments of varying lengths, from 1 to 30 s prior to question onset. Each window incorporated the most recent activity but varied in the amount of past data included, based on the assumption that more recent activity is more predictive of behavior. Testing multiple window lengths allowed us to examine which temporal scale provided the most robust information about creativity potential. This procedure yielded 25,650 single-trial coherence features (171 connections × 5 frequency bands × 30 windows) to predict the scalar value of regularized *C*.

**Figure 2.**
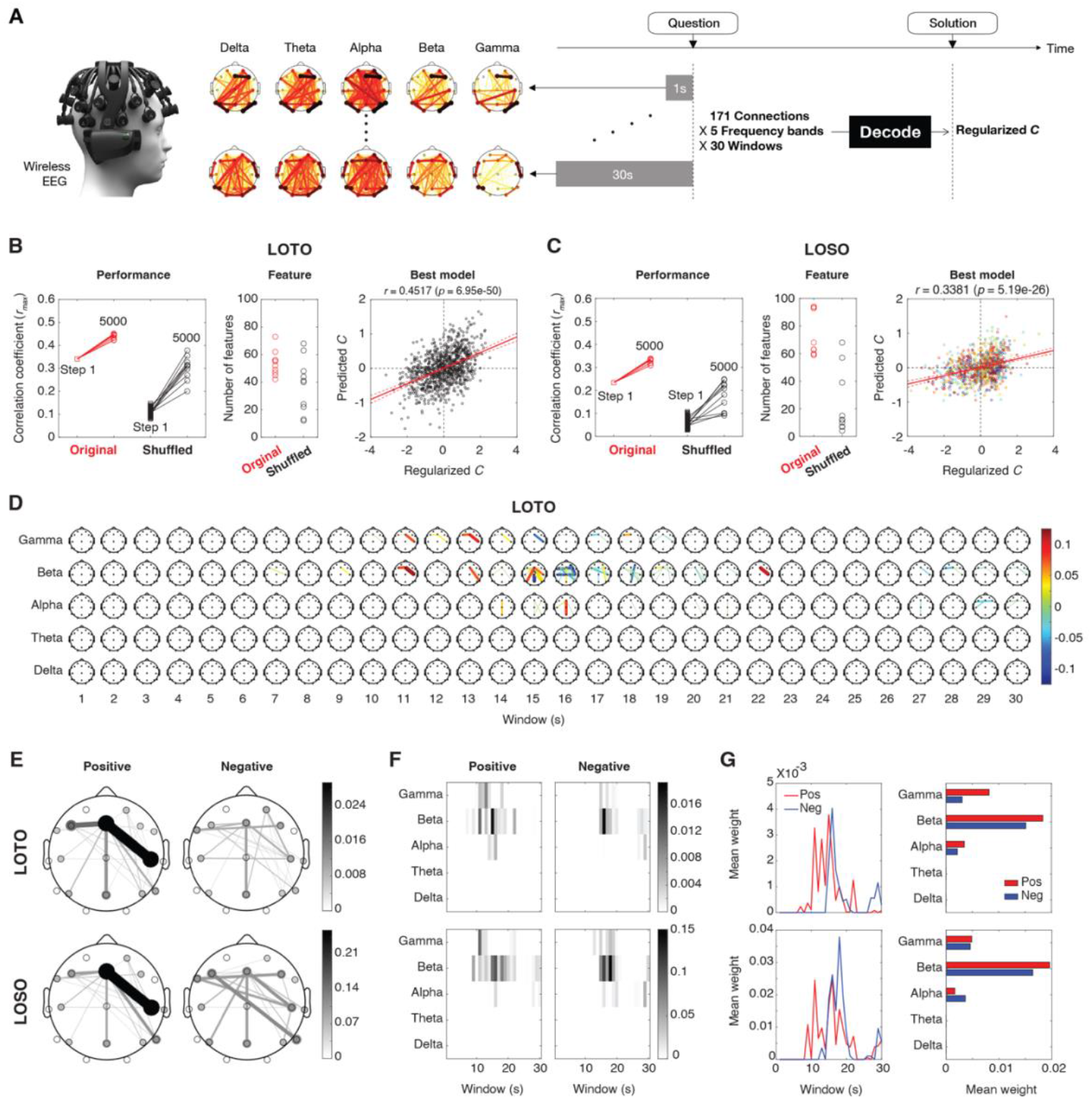
Decoding performance and model characteristics. (**A**) Decoding strategy. A trial timeline is shown at the top right. Resting-state EEG segments of varying lengths (1∼30 s before question onset) were analyzed to compute coherence across five frequency bands, yielding single-trial EEG features (171 connections × 5 frequency bands × 30 windows). These features were used to predict the corresponding single-trial value of regularized *C*. (**B**) Decoding performance with leave-one-trial-out (LOTO). Left: Performance was measured by the maximal correlation coefficient (*r*_*max*_) at Step = 1 (square) and Step = 5000 (circle) for both the original and shuffled data. For Step = 1, 500 sets of shuffled data were generated, but only 10 sets were decoded, with one set decoded only once. In contrast, the original data consisted of one dataset, but 10 runs were performed, each with random pruning. Middle: The number of selected features after 5000 steps is shown on the right for both original and shuffled runs. Right: Best model. The model with the highest correlation coefficient among the 10 runs is displayed. Regularized and predicted *C* values are plotted for each trial, with the correlation coefficient and corresponding p-value indicated. Red line: linear regression; dashed lines: 95% confidence interval. The same plots, color-coded by question index and subject index, are shown on the right. (**C**) Decoding performance with leave-one-subject-out (LOSO). The same representation as in panel B is used. In the right panel, each trial in the best model is color-coded by subject index. (**D**) Feature weights. Weights of all features averaged across 10 models with LOTO are shown, where the color and thickness of each connection reflect the weight magnitude. (**E**) Summary of spatial distributions of absolute feature weights for LOTO and LOSO, with positive and negative weights shown separately. (**F**) Summary of spectrotemporal distributions for positive and negative weights for LOTO (top) and LOSO (bottom). (**G**) Summary of temporal and spectral distributions for positive and negative weights for LOTO (top) and LOSO (bottom). The EEG image in panel A is from the Brain Products website.

We applied partial least squares regression (PLSR) with a leave-one-trial-out (LOTO) cross-validation scheme. Decoding began with the top 200 features showing the highest correlation with regularized *C* at Step = 1. Features were then iteratively pruned to optimize decoding performance, defined as the maximal correlation coefficient (*r*_*max*_) across different numbers of PLS components (see Figure S1 for analytical pipeline and Methods for details). The decoding process was terminated at Step = 5000. As a control, we randomly shuffled the regularized *C* values across trials 500 times and repeated the same decoding procedure.

At Step = 1, the original data (top 200 features) achieved a decoding performance of *r* = 0.341, whereas shuffled data yielded *r* = 0.11 ± 0.01 (mean ± standard deviation), which was significantly lower (*p* ≈ 0, one-tailed t-test; Figure 2B, left). For the original data, we ran 10 independent decoding iterations with different random pruning sequences. By Step = 5000, decoding performance improved to 0.44 ± 0.01, with a maximum of 0.45. For the shuffled data, 10 datasets were each decoded once, yielding performance of 0.30 ± 0.05, still significantly below the original data (*p* = 4.66e-6). The number of features retained after 5000 steps decreased from 200 to 53.7 ± 9.0 for the original data and to 37.5 ± 19.5 for the shuffled data (Figure 2B, middle). The correlation between regularized and predicted *C* in the best-performing model (*r* = 0.45, *p* = 7.0e-50) is shown in Figure 2B (right).

We also performed decoding using a leave-one-subject-out (LOSO) cross-validation scheme (Figure 2C). The model achieved a maximal performance of *r* = 0.34 (*p* = 5.2e-26), which was significantly higher than the shuffled baseline (*p* = 1.59e-05). To examine whether this correlation was driven by individual subject differences (e.g., consistently higher or lower creativity scores), we plotted the results with trials color-coded by question index and subject index, which confirmed that the observed correlation was not attributable to such biases (Figure 2C, right). This demonstrates that the decoding model can generalize to new subjects, making it more practical for real-world applications where model training and behavioral data collection are often difficult (see Discussion).

### Creativity potential network (CPN): decoding model

We next examined the feature weights of the decoding model. The weights averaged across 10 models (using LOTO as an example) are shown in Figure 2D, which we refer to as the creativity potential network (CPN). To visualize spatial distributions, we separated positive and negative weights and averaged them across frequency bands and time windows, revealing distinct positive and negative networks for both LOTO and LOSO (Figure 2E). The results show similar patterns for LOTO and LOSO, with enhanced interactions between frontal and right temporal regions associated with higher creativity potential.

We further visualized spectrotemporal patterns by averaging positive and negative weights across connections (Figure 2F), and then across frequency bands and time windows to highlight temporal and spectral contributions (Figure 2G). The results were consistent between LOTO and LOSO, showing that ∼15 s windows contributed most strongly for both positive and negative weights. Suppressive networks engaged earlier (∼15 s before question onset), followed by enhanced networks (∼10 s before onset). Contributions were centered primarily in the beta band.

### Monitoring creativity potential

We next applied the decoding model to monitor creativity potential continuously. Several 5-min resting sessions were recorded prior to the experiments. By applying the LOTO decoding model to these resting-state data, we tracked the predicted *C* (creativity potential) over time. As expected, predicted *C* fluctuated, sometimes with slower drifts (see examples in Figure 3A). After excluding sessions with missing values caused by jump noise and eye-movement artifacts during EEG preprocessing (see Methods), we analyzed 32 complete, artifact-free sessions from 16 subjects (Figure 3B) and calculated the dominant period of each time course (Figure 3C). The median and mean periods were 3.4 and 3.2 min, respectively, indicating that creativity potential cycled approximately every 3 minutes during rest.

**Figure 3.**
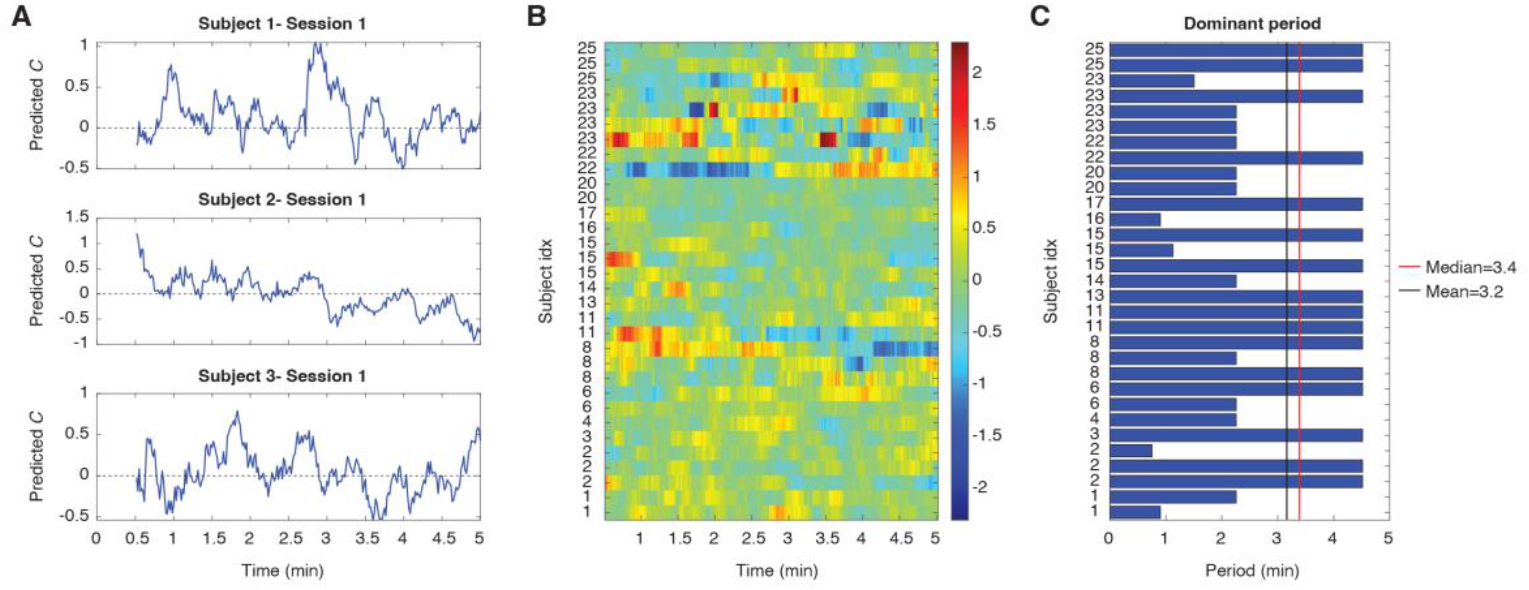
Dynamics of creativity potential. (**A**) Examples of predicted *C* values during 5-min resting sessions from three subjects. Note that the first prediction is available only after 30 s of data were collected. (**B**) Predicted *C* values across all sessions from 16 subjects, with color indicating the magnitude of predicted *C*. (**C**) Dominant periods extracted from the time courses of each session. Both the median and mean values are shown.

## Discussion

Creativity is a dynamic process shaped not only by domain knowledge and motivation but also by the brain’s intrinsic state. Here, we identified a neural marker of creativity potential, the preparatory brain activity that predicts whether upcoming problem solving will be more or less creative. This marker generalized across two complementary tasks, AUT and FIT, as well as across individuals, suggesting that it reflects a shared, domain-general capacity rather than task-specific strategies. Moreover, creativity potential fluctuated in intrinsic cycles of approximately three minutes, pointing to endogenous brain rhythms that may constrain the temporal dynamics of creative thought.

These findings extend prior work linking creativity to network flexibility and large-scale neural dynamics. Converging evidence suggests that creative cognition depends on the flexible reconfiguration of brain networks, particularly the interplay between the default mode network, executive control network, and salience network (1, 5, 16–18). In particular, frontal-temporal interactions have been implicated in semantic integration and remote association (17, 19), processes critical for generating novel ideas. Our decoding model revealed a CPN characterized by enhanced frontal-temporal coupling prior to creative performance, consistent with these earlier observations. Temporally, suppressive networks engaged earlier, followed by enhanced networks closer to problem onset, echoing prior EEG findings showing preparatory neural activity as a determinant of creative outcomes (6, 20, 21).

These results also resonate with predictive coding accounts of brain function, which propose that perception and cognition emerge from the interplay between top-down predictions and bottom-up prediction errors (22, 23). From this perspective, creativity may reflect moments when predictive models are loosened, allowing the exploration of less expected possibilities before reconverging on effective solutions (24). The transient disengagement of suppressive networks observed here may correspond to reduced precision-weighting of priors, thereby enabling exploration of more diverse solution spaces. Conversely, the subsequent enhancement of frontal–temporal networks may support integration and selection, consistent with the need to balance divergent and convergent thinking (25, 26).

A novel contribution of this study is the discovery that creativity potential fluctuates in intrinsic cycles of approximately three minutes during rest. Spontaneous fluctuations in attention, vigilance, and mind-wandering are well documented (27, 28), and our findings suggest that creativity potential is similarly dynamic. The ∼3-minute periodicity may relate to ultradian rhythms in neural, behavioral, and physiological systems, which provide functional timescales for alternating between exploration and exploitation modes of cognition (29). Such intrinsic oscillations may help explain why creative insights often occur unpredictably and why creativity is difficult to sustain continuously over time.

Methodologically, this work advances the study of creativity in several ways. First, by combining divergent and convergent thinking tasks with automated GPT-based evaluation, we provide a scalable and multidimensional measure of creativity that reduces reliance on subjective judgment (10, 11). Second, by focusing on preparatory EEG activity, we establish a neural marker that predicts creative outcomes in advance, complementing prior work that has largely examined activity during or after solution generation (17, 20). Third, by applying the decoding model to resting-state data, we demonstrate that creativity potential can be monitored continuously, opening possibilities for real-world applications. These include neurofeedback systems that allow individuals to train brain flexibility (30, 31) and adaptive environments that time interventions to periods of high creativity potential.

Together, these findings provide converging evidence for a domain-general neural marker of creativity potential, grounded in flexible brain network dynamics. By situating creativity within predictive coding theory and revealing its intrinsic temporal cycles, this study bridges cognitive neuroscience and practical applications, offering new avenues for monitoring and fostering creativity in everyday life.

## Methods

### Participants

Data were collected from 28 participants (17 males, 11 females), aged 18∼29 years (mean ± standard deviation: 22 ± 2.5 years). All were university students, native Japanese speakers, with no history of neurological disorders. Prior to the experiment, participants were informed of the procedures and provided written informed consent. The study protocol and consent form were approved by the Research Ethics Committee of the University of Tokyo (No. 23-27). Participants were recruited through an online advertisement (https://www.jikken-baito.com).

### Experimental setup

Data were collected in a quiet, well-lit room. Participants sat in front of a 43-inch monitor for the presentation of questions and instructions and responded using a separate tablet with a keyboard. EEG signals were recorded with the CGX Quick-20r v2 wireless dry-electrode system (Cognionics, Inc., San Diego, CA), comprising 20 channels (including a reference electrode at the left earlobe) arranged according to the international 10–20 system. Signals were sampled at 500 Hz, and electrode impedance was maintained below 2500 kΩ throughout recording.

### Experimental procedure

Each participant completed 30 AUT questions and 30 FIT questions, divided equally across two days. On each day, participants performed one block of 15 AUT questions and one block of 15 FIT questions. The order of blocks was reversed on the second day, and the starting task (AUT or FIT) was counterbalanced across participants. After completing both experimental days, participants filled out online questionnaires assessing their everyday creative behavior.

At the start of each session, participants were fitted with the dry EEG system. A five-minute resting-state recording was then conducted, during which participants sat still, kept their eyes open, and fixated on a central cross. Each task block began with one practice trial followed by 15 experimental trials. Between blocks, there was a five-minute break; the EEG cap was removed, and participants were free to walk or exercise. The same procedures were repeated on the second day, with only the order of task blocks changed.

All stimuli were presented on a black background. At the beginning of each block, participants read on-screen instructions and pressed the SPACE key to start. Each trial began with a fixation cross presented for 30 s (used for subsequent decoding analyses), during which participants were instructed to remain still and maintain fixation, as in the resting-state recording. A question then appeared, and participants were asked to provide a single answer as quickly as possible. They pressed the SPACE key when ready, typed their response into a Google Form on a tablet, and pressed SPACE again to proceed. Response time was defined as the interval between question onset and the keypress. Participants could take short breaks during inter-trial intervals.

Each trial included a countdown timer above the question: 90 s for AUT and 180 s for FIT. If participants failed to press SPACE before the timer expired, the trial was marked as unanswered. If they began responding before time ran out, they were allowed to finish typing their answer even if it exceeded the time limit.

### Response evaluation and regularization

Participants’ responses were evaluated automatically using GPT-4o, with prompts adapted from previous studies (AUT: Kern et al., 2024; FIT: Wu et al., 2024). For AUT, each response was rated on two dimensions: Novelty and Feasibility. For FIT, each response was rated on three dimensions: Novelty of item combinations, Feasibility of item combinations, and Goal Attainment. All scores were scaled from 1 to 100. For AUT, overall creativity (AUT-*C*) was defined as the geometric mean of *N* and *F* (*C* = √(*N* × *F*)). For FIT, overall creativity (FIT-*C*) was defined as the geometric mean of *N, F*, and *G* (*C* = ^3^√(*N* × *F* × *G*)).

To account for differences in scoring distributions and response tendencies, we applied a two-level regularization procedure on AUT-*C* and FIT-*C*. First, scores were z-scored within each question to normalize performance relative to that item’s difficulty. This step reduces potential bias from certain questions eliciting systematically higher or lower ratings. Second, the question-normalized scores were z-scored again within each participant, thereby controlling for inter-individual response biases (e.g., consistently high-or low-scoring participants). This double-regularization ensured that creativity scores reflected relative performance both within and across participants, while minimizing the influence of task-specific or individual-level scaling differences.

### Creativity questionnaires

After completing the 2-day task, participants filled out four standardized creativity questionnaires:

1. Kaufman Domains of Creativity Scale (KDOCS) (12): Assesses self-perceived creativity across five domains: everyday, scholarly, performance, mechanical/scientific, and artistic. Participants rate themselves on a 5-point Likert scale from 1 (much less creative) to 5 (much more creative).
2. Creative Achievement Questionnaire (CAQ) (13): Measures real-world creative accomplishments across multiple domains, ranging from “no training” to “national recognition/prizes.”
3. Runco Ideational Behavior Scale (RIBS) (14): A 23-item scale assessing creative potential through the frequency of idea generation and expression. Responses are given on a 5-point Likert scale from 1 (never) to 5 (almost every day/more than twice a day).
4. Inventory of Creative Activities and Achievements (ICAA) (15): Captures both creative activities and achievements across multiple domains. For each domain, it records (a) the frequency of engagement in creative activities and (b) the highest level of achievement. Compared with the CAQ, the ICAA provides a more balanced assessment by including both formal accomplishments and informal activities.

For each subject and questionnaire, subcategory scores were summed to produce a single scalar value representing the overall score for that questionnaire.

### EEG preprocessing

EEG was recorded from 20 electrodes and analyzed on 19 channels (excluding the reference). Preprocessing was conducted with the MATLAB FieldTrip toolbox (32). For each trial, we analyzed the pre-trial interval immediately preceding question onset. Data were downsampled to 125 Hz, high-pass filtered at 0.2 Hz, and line noise was removed at 50 and 100 Hz. Signals were then re-referenced using a common-median reference (ft_preprocessing.m). Automatic artifact detection targeted jump noise (ft_artifact_jump.m) and electrooculogram (EOG) artifacts (ft_artifact_eog.m), with contaminated trials excluded using complete rejection (ft_rejectartifact.m).

### Connectivity analysis

Spectral power and cross-spectral density were estimated with multitaper FFT (frequency smoothing = 5 Hz; 0.2–40 Hz; ft_freqanalysis.m). Pairwise coherence was computed (ft_connectivityanalysis.m) and averaged across five canonical frequency bands: delta (0.1– 3.5 Hz), theta (3.5–8 Hz), alpha (8–12 Hz), beta (13–30 Hz), and gamma (30–40 Hz). This procedure was repeated over pre-trial windows immediately before onset (e.g., 1–30 s), yielding a trial-wise tensor of 171 connections × 5 frequency bands × 30 windows.

### Decoding analysis (LOTO)

For each connection, frequency band, and window, trial-wise coherence values were correlated with single-trial regularized *C* scores across *K* − 1 training trials (*K*=168, combining AUT and FIT). Features were ranked by their correlation coefficients, and the top *N* = 200 features were retained for decoding using PLSR. Note that the regularization of *C* was also applied within each set of *K* − 1 training trials. The overall pipeline is illustrated in Figure S1.

PLSR was performed with the number of components ranging from 1 to *M* = 5 (plsregress.m). We limited the model to five components because adding more substantially increased computation time without yielding clear performance gains. In LOTO cross-validation, each held-out trial produced *K* × *M* correlation coefficients, reflecting the match between predicted and actual *C*. For each validation, we selected the maximum correlation across components as the trial’s decoding performance. Averaging these values across all trials yielded the overall decoding performance metric (*r*_*max*_) for that iteration.

To further improve performance and reduce dimensionality, we implemented a greedy feature pruning heuristic. At each iteration, one active feature was randomly removed; if this removal increased *r*_*max*_, the change was kept, otherwise the algorithm retried with another feature. If no improvement was observed after 50 attempts, a random inactive feature was added back into the model. This iterative pruning process continued until 5,000 steps were completed, yielding a compact and optimized feature set for decoding.

### Decoding analysis (LOSO)

Similar to LOTO approach, LOSO decoding began by correlating trial-wise coherence values with single-trial regularized *C* scores across the training set. The key difference is that, in LOSO, all trials from one subject (60 trials) were excluded from training and used as the test set. The model was then trained on the remaining *K* – 60 trials and evaluated on the held-out subject.

### Monitoring CPN during 5-min resting

The same EEG preprocessing pipeline was applied to resting-state data. Each 5-min session was segmented into 30-s windows using a 1-s moving step. For each segment, connectivity analysis was performed, yielding a feature matrix of 171 connections × 5 frequency bands × 30 windows. These features were then input into the best LOTO decoding model (Figure 2D) to generate predicted *C* scores. The first prediction was obtained at 30 s, with subsequent predictions updated every 1 s. Sessions containing excessive jump noise or EOG artifacts were excluded for simplicity. After artifact rejection, the dataset was reduced from 57 sessions across 28 subjects to 32 sessions across 16 subjects.

## Author contributions

Z.C.C. conceptualized the study. T.L.L and M.S. implemented the experiment and collected the data. Z.C.C. performed data analysis and wrote the paper. T.L.L and M.S. edited the paper. All authors contributed to and approved the final paper.

## Acknowledgments

We thank Yansen Wang (Microsoft Research Asia) for his suggestions on the decoding analysis, and Dr. Yu-Shian Su for acquiring the GPT scores. This study was supported by the World Premier International Research Center Initiative (WPI), MEXT, Japan (to Z.C.C.), and the IRCN–Daikin SCP (to Z.C.C.).

## Supplemental figures

**Figure S1.**
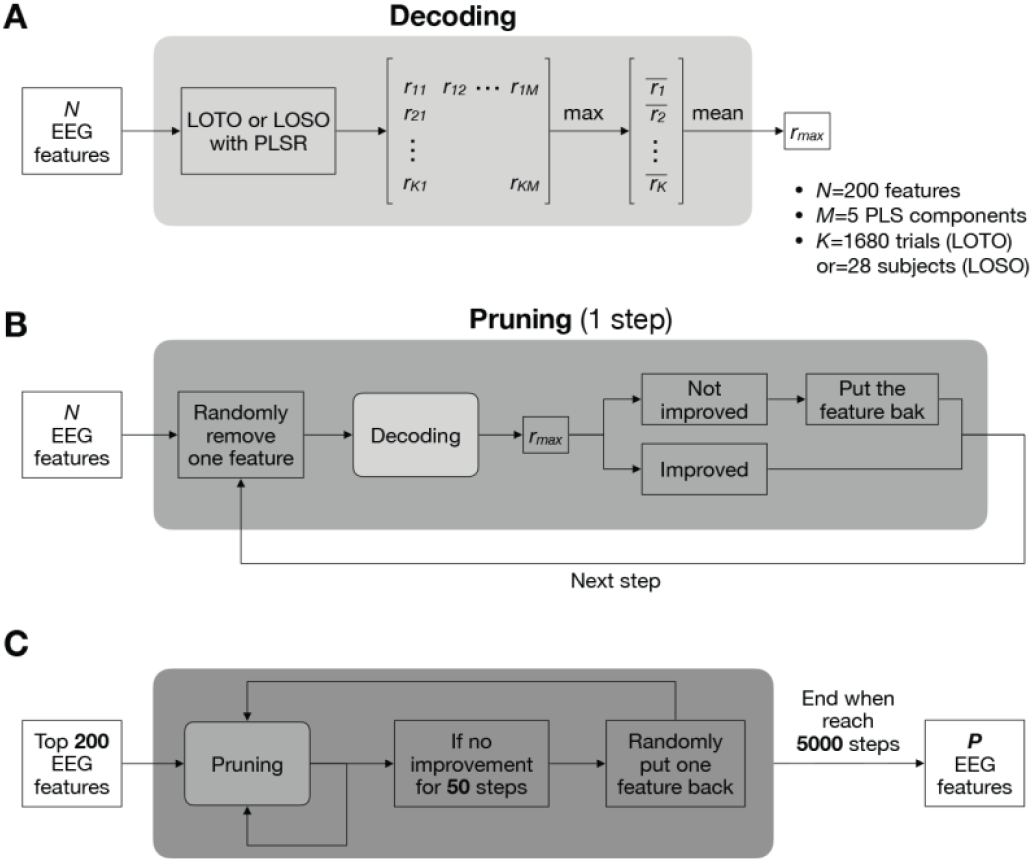
Decoding pipeline. (**A**) Decoding procedure. The top *N* (=200) EEG features were used to predict regularized *C*. Using leave-one-trial-out (LOTO) cross-validation with partial least squares regression (PLSR), *K* (=1680) correlation coefficients (*r*) were obtained for each partial least squares (PLS) component (*M* = 5). For each left-out trial, the maximal *r* across components was selected, and the average across trials was taken as the decoding performance (*r*_*max*_). For leave-one-subject-out (LOSO) decoding, trials were replaced with subjects (*K* = 28). (**B**) Feature pruning (single step). Illustration of the pruning process applied in one iteration. (**C**) Feature pruning and reintroduction. Across 5000 steps, the number of features was reduced from 200 to *P*, with some pruned features potentially added back to improve the performance.

